# Dorsal Premotor Contributions to Auditory Rhythm Perception: Causal Transcranial Magnetic Stimulation Studies of Interval, Tempo, and Phase

**DOI:** 10.1101/368597

**Authors:** Jessica Ross, John Iversen, Ramesh Balasubramaniam

**Affiliations:** University of California, Merced; University of California, San Diego

**Author notes:** Address for correspondence: Jessica M. Ross, Cognitive & Information Sciences, University of California, Merced, CA 95343.

## Abstract

It has been suggested that movement planning networks are critical for time perception. The Action Simulation for Auditory Prediction (ASAP) hypothesis proposes that the dorsal auditory stream is involved in predictive beat-based timing through bidirectional interchange between auditory perception and dorsal premotor (dPMC) prediction via parietal regions, as has been supported by brain imaging and transcranial magnetic stimulation (TMS). However, causal impact of dPMC on time perception has not been tested directly. We used a TMS protocol that down-regulates cortical activity, continuous theta burst stimulation (cTBS), to test for causal contributions of left dPMC to time perception. Three experiments measured (1) discrete interval timing perception, and relative beat-based musical timing for (2) tempo perception and (3) phase perception. Perceptual acuity was tested pre- and post-cTBS using a test of sub-second interval discrimination and the Adaptive Beat Alignment Test (A-BAT). We show (N = 30) that cTBS down-regulation of left dPMC interferes with interval timing perception and the ability to detect differences in musical tempo, but not phase. Our data support causal involvement of premotor networks in perceptual timing, supporting a causal role of the left dPMC in accurate interval and musical tempo perception, possibly via dorsal stream interactions with auditory cortex.

## Introduction

Perception of musical beat is a predictive form of time perception, distinct from absolute interval timing^1–5^. In absolute timing, intervals are perceived and encoded discretely, but in relative, beat-based timing, intervals are interpreted relative to a perceived and ongoing beat structure^5^. Beat perception entails predictions about tempo (beat period) and about phase (beat onset times). The mechanisms involved with making beat-based timing predictions are of interest for a number of reasons. One reason is that beat perception seems to be a human ability, with only minimal analogues in some non-human species. Beat perception is also a defined test case for the study of sensorimotor interactions, with a rich literature on prediction and error correction in finger-tapping synchronization to auditory rhythms^6,7^. Recent evidence suggests that in some scenarios the motor system might actively shape the perception of sound^2,3,8^.

The dorsal premotor cortex (dPMC) is involved with movement planning, including for sound guided motor synchronization^9,10^. Additionally, dPMC is active during purely perceptual timing tasks, in the absence of overt movement^11,12^. However, it remains unknown if such ‘purely perceptual’ activity in dPMC is necessary to perceptual processing of the beat, or if it is an epiphenomenal consequence of planning for unexpressed overt movement^13^. A number of accounts have hypothesized an active motor role in perception^2,8^. One, which predicts a specific role for dPMC, is the ASAP hypothesis, which proposes that motor planning regions are involved in making beat-based timing predictions that are causally necessary for beat perception^3^. Further, the dorsal auditory pathway is hypothesized to be where auditory and motor networks interact to compare timing predictions in motor cortex with incoming sounds^3^. The dorsal auditory pathway connects caudal auditory regions, such as posterior superior temporal gyrus, with dorsal frontal premotor regions, such as dorsal premotor cortex, via parietal regions such as the angular gyrus, and this pathway is bi-directional^14^. dPMC is part of the dorsal auditory stream, so finding its involvement in beat-based timing would support the ASAP hypothesis.

ASAP posits that beat-based timing relies on internal predictive models that are continuously updated^1,6,7^, and describes how some aspects of beat perception support that an internal predictive model is being used^3^. These aspects include negative mean asynchrony which is thought to demonstrate timing prediction^7,15–17^, and tempo flexibility in the perception of rhythmic structure^18–21^, the susceptibility of beat perception to willful control, and improved perceptual acuity of events that occur on the beat^22^, which all support that top-down predictions can influence auditory perception. In addition to this evidence, beat perception has been shown to be directly influenced by body movement^23–28^, which supports that motor behavior or planning may also influence auditory perception^2,3^. In support of the proposal that beat perception uses the dorsal auditory stream, we show in previous work that TMS-induced down-regulation of posterior parietal cortex, a critical link between premotor and auditory regions in this pathway, interferes with phase aspects of beat perception^29^.

The dorsal auditory pathway, also referred to as the dorsal stream, is associated with localization of sounds in space, phonological processing and sensorimotor integration and control of speech^14,30–32^. The dorsal stream includes both afferent and efferent tracts, enabling bidirectional communication between auditory and premotor cortex. There is some evidence that the dorsal stream is involved in auditory temporal processing^33^, with a suggested role in musical processing: in imagined time-reversed musical melodies^33,34^ and in musical phase perception^29^.

However, a central question that has not yet been directly tested is if dPMC has a causal role in timing and beat perception. One method to directly assess causal contributions is through transient modulation of cortical function using transcranial magnetic stimulation (TMS). Specifically, TMS can be used to determine causal contributions to perceptual tasks of different brain areas by functionally modulating cortical excitability and observing changes in perception. Continuous theta burst stimulation (cTBS), a TMS protocol, down-regulates focal cortical excitability through hyperpolarization of cell bodies, which can be measured behaviorally^35^.

Two prior studies have used brain stimulation to probe brain mechanisms for different aspects of timing behavior. Grube et al. (2010, see ref. 36) used cTBS to demonstrate causal involvement of motor networks in absolute interval timing perception and Pollok et al. (2017, see ref. 37) used a related technique, tDCS, to demonstrate causal contributions of dorsal premotor regions to rhythm reproduction using an auditory-motor synchronization and continuation tapping paradigm^37^. In previous work using cTBS, we show that down-regulation of left PPC interferes with accurate perception of beat phase timing in music but not beat tempo timing or single absolute interval discrimination^29^. Notably, this work examined changes in timing perception using tasks that did not require explicit motor synchronization. As PPC is an intermediary node in the dorsal auditory stream, mediating between auditory and premotor cortices, this provided support for the ASAP hypothesis. The current study tests the next link in the chain as, to our knowledge, brain stimulation techniques have not been used previously to observe changes in timing perception without motor synchronization with down-regulation of dPMC.

To explore specific causal contributions of dPMC to timing, we present three targeted studies to address the following questions: (1) Is dPMC causally involved in absolute interval timing perception? (2) Is dPMC causally involved in musical beat tempo timing perception? (3) Is dPMC causally involved in musical beat phase timing perception? For this work, we targeted left dPMC (Figure 1, Talairach −32, −12, 62; coordinates taken from ref. 38) and tested timing perception performance before and after cTBS. We contrasted this with timing perception performance before and after a sham cTBS stimulation, in which participants believed they were receiving brain stimulation but were not. Based on Pollok et al. (2017, see ref. 37), we might have expected dPMC to be involved in both absolute and relative timing, but because they used a continuation tapping task, we were unsure whether this role of dPMC for internally generated rhythm would be sufficient to support predictions about absolute interval timing perception, and therefore were unsure whether to expect cTBS down-regulation of left dPMC to disrupt accurate interval timing perception. Based on the proposals set forth by the ASAP hypothesis^3^, we expected to find evidence that dPMC is causally involved in both tempo and phase timing.

**Figure 1.**
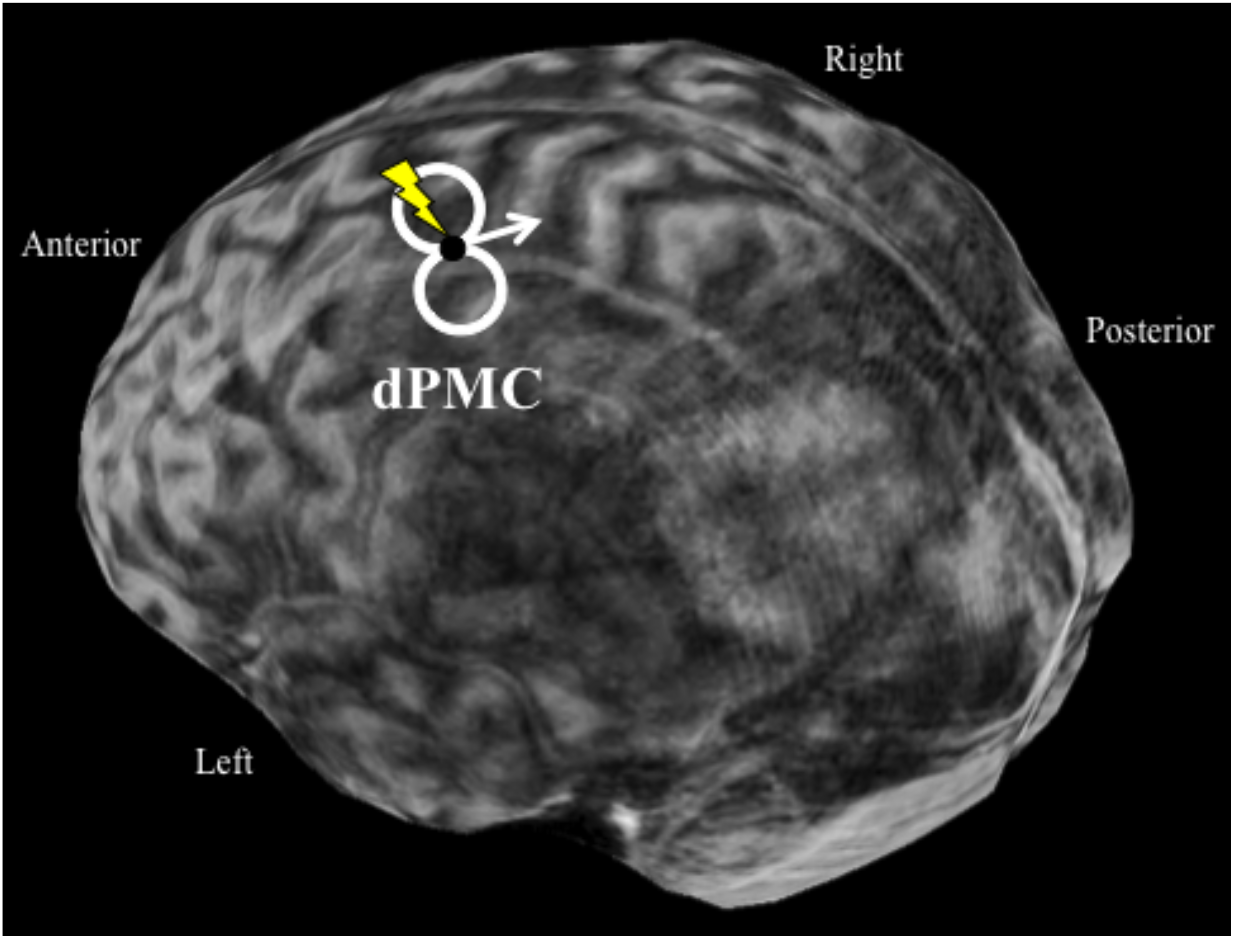
Left dorsal premotor cortex stimulation target and coil orientation. Center of coil was placed at Talairach −32, −12, 62^38^, with the coil facing anteriorly to induce an anterior to posterior flow of current (indicated here with an arrow)^46^.

## Results

Analysis of pre- to post-cTBS changes in perceptual acuity was completed for each timing experiment (See Materials and Methods below and Figure 2 for more details on the perceptual tests), with N=30 completing all three tests, using paired samples t-tests. Additionally, to support t-tests and probe robustness of the findings, pre- to post-cTBS changes were compared across sham and left dPMC stimulation conditions using linear mixed effects models, with a fixed effect for pre- versus post-cTBS and random effects for condition and for participant. P values were obtained by likelihood ratio tests of the full model with the effect in question against the model without the effect in question^39^.

**Figure 2.**
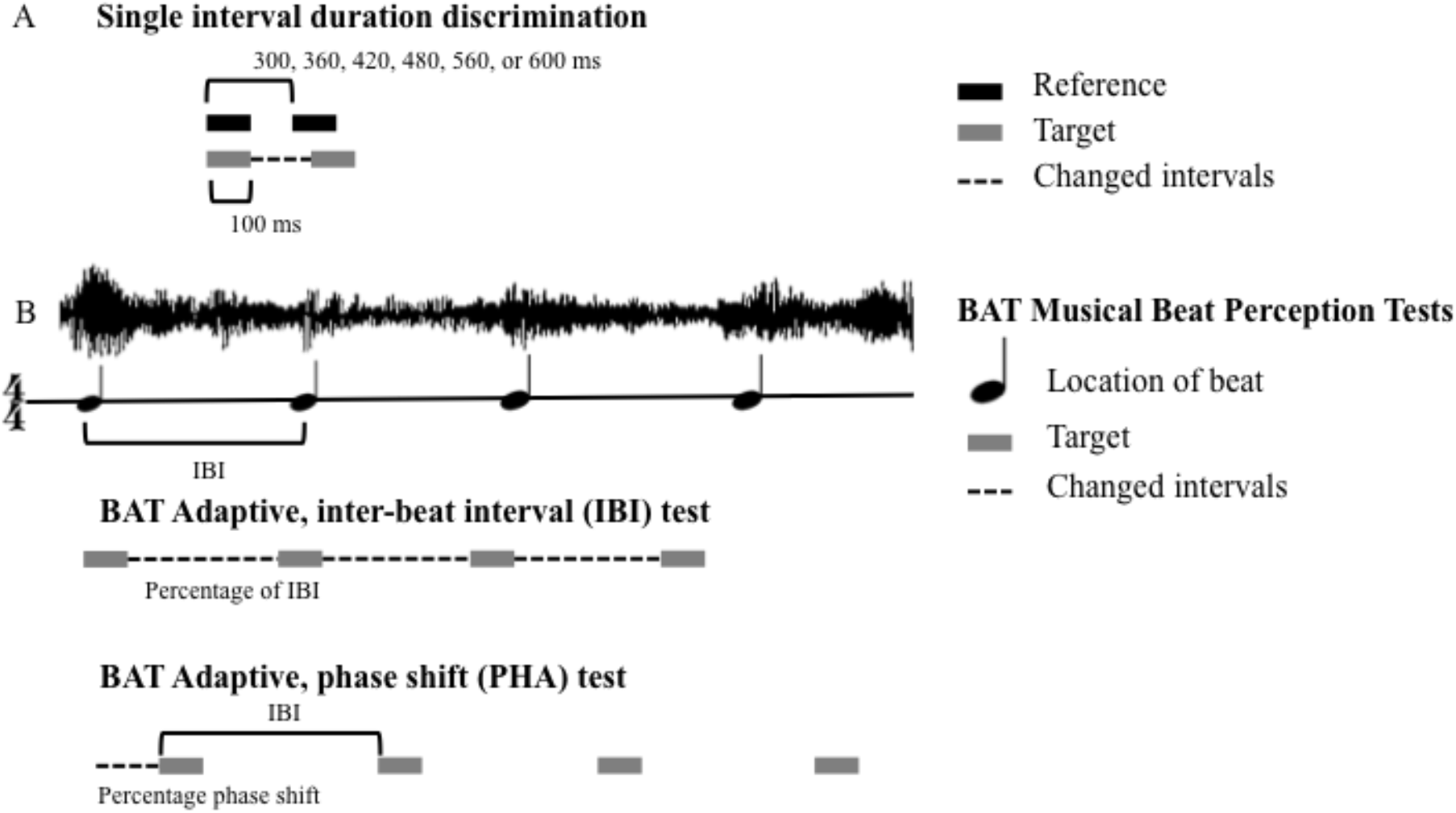
Adaptive auditory timing tests used for determining perceptual thresholds. (A) Experiment 1: Single-interval duration discrimination test^29,36^ (B) Experiment 2/3: Tests of musical timing perception (A-BAT)^29,45^, used to determine perceptual thresholds for detecting musical tempo (experiment 2) and phase alignment (experiment 3).

### Interval Timing Discrimination

T-test comparison of interval discrimination thresholds pre- to post-cTBS was significant, with a 22.87% higher threshold after the treatment (interval difference threshold: pre = 48.22 ± 6.88%, post = 59.25 ± 8.56%; t(29) = −2.083, p = .046, Cohen’s d = 0.38). This higher threshold after down-regulation of left dPMC indicates a decrease in perceptual acuity for differentiating single interval durations. As a control, we found no pre- to post-cTBS difference with sham stimulation (interval difference threshold: pre = 35.39 ± 5.00%, post = 37.84 ± 4.57%; t(29) = −0.606, p = .549, Cohen’s d = 0.11; See Figure 2A for more on the interval timing test; See Figure 3A for interval thresholds). A linear mixed effects model revealed no significant changes across sham and dPMC conditions for pre- to post-cTBS down-regulation (χ^2^ (1) = 2.3116, p = .1284), meaning this model does not support an effect of stimulation site on the pre- to post-cTBS threshold change.

**Figure 3.**
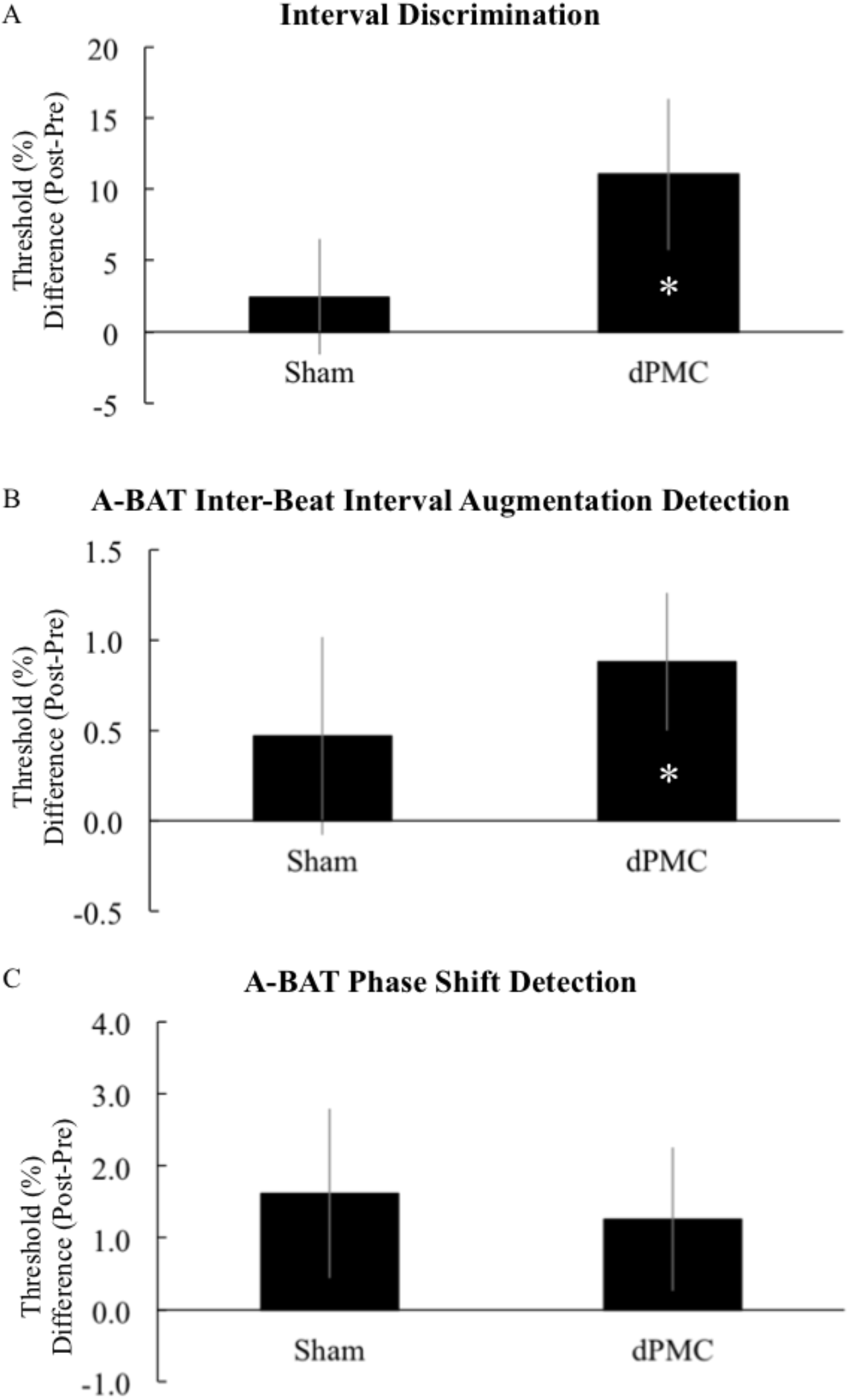
Mean post-cTBS minus pre-cTBS threshold differences for the three timing perception experiments in the left dorsal premotor stimulation condition and sham stimulation. Error bars represent ±1 standard error from the mean. Asterisks indicate significance at p < .05 (A) Experiment 1: Single-interval duration discrimination^29,36^ (B) Experiment 2: Musical tempo detection (A-BAT IBI)^29,45^ (C) Experiment 3: Musical phase detection (A-BAT Phase)^29,45^. There was an increase in detection thresholds pre- to post-cTBS in experiment 1 (t(29) = –2.083, p = .046, Cohen’s d = 0.38) and experiment 2 (t(29) = −2.318, p = .028, Cohen’s d = .42) with left dPMC down-regulation.

### Tempo Timing Perception

T-test comparison of tempo detection thresholds pre- to post-cTBS was significant, with a 19.93% higher threshold after the treatment (tempo deviation threshold: pre = 4.41 ± .52%, post = 5.29 ± .57%; t(29) = −2.318, p = .028, Cohen’s d = .42). This higher threshold after down-regulation of left dPMC indicates a decrease in perceptual acuity for detecting differences in tempo. As a control, we found no pre- to post-cTBS difference with sham stimulation (tempo deviation threshold: pre = 4.59 ± .54%, post = 5.06 ± .48%; t(29) = -.857, p = .399, Cohen’s d = 0.16; See Figure 2B for more on the tempo timing test; See Figure 3B for tempo detection thresholds). A linear mixed effects model revealed no significant changes across conditions for pre- to post-cTBS down-regulation (χ2 (1) = 3.3225, p = .0683), meaning this model does not support an effect of stimulation site on the pre- to post-cTBS threshold change.

### Phase Timing Perception

T-test comparison of phase detection thresholds pre- to post-cTBS was not significant (phase shift threshold: pre = 15.81 ± 1.55%, post = 17.07 ± 1.59%; t(29) = −1.265, p = .216, Cohen’s d = .23), thus the data do not support a change in perceptual acuity for detecting changes in phase. We also found no pre- to post-cTBS difference with sham stimulation (phase shift threshold: pre = 15.36 ± 1.33%, post = 16.98 ± 1.36%; t(29) = −1.375, p = .180, Cohen’s d = 0.25; See Figure 2B for more on the phase timing test; See Figure 3C for phase detection thresholds). A linear mixed effects model revealed no significant changes across conditions for pre- to post-cTBS down-regulation (χ2 (1) = 3.2291, p = .0723), meaning this model does not support an effect of stimulation site on the pre- to post-cTBS threshold change.

## Discussion

Using focal down-regulation of left dPMC with cTBS brain stimulation, the present series of experiments tested for specific causal roles of left dPMC in different aspects of timing perception. These experiments were designed to observe interval timing perception^29,36^ (Figure 2A), musical tempo perception and musical phase perception^29^ (Figure 2B). We found (N = 30) that cTBS down-regulation of left dPMC interferes with two aspects of timing perception: interval timing perception acuity (Figure 3A) and the ability to detect changes in musical tempo (Figure 3B).

In a design similar to the one presented here, Grube et al. (2010, see ref. 36) used cTBS to demonstrate causal involvement of motor networks in timing perception. In their work, cTBS to medial cerebellum raised perceptual thresholds on a test of absolute interval timing perception, but not on a beat-based timing test. Their work supports that networks used for absolute interval timing and for relative, beat-based timing are distinct. The work presented here also supports that absolute interval timing perception may rely on different networks than some forms of beat-based timing (here phase timing perception), but perhaps not on others (here tempo perception). We show that aspects of beat perception appear to be separable, but also that interval timing perception and beat-based timing may have points of overlap.

Interestingly, and contrary to our hypothesis, while our past cTBS study found that PPC down-regulation affected phase perception, with the dPMC target we did not find any evidence of disruption of musical phase timing (Figure 3C). Our initial hypotheses were based on the, possibly simplistic, notion that the entire dorsal stream would be equally involved in all aspects of beat timing, such that disruption of any node in the stream would lead to both tempo and phase timing effects. However, the pattern of results across our two studies suggest that tempo and phase timing might reflect distinct timing mechanisms sub-served by different nodes or networks^6,7^ or different motor network hubs^37^. Numerous prior behavioral studies have suggested that in sensorimotor synchronization, tempo and phase may be supported by distinct processes, evidenced, for example, by the differences in tempo and phase error correction in sensorimotor synchronization^6,7^. Repp (2005, ref. 7) suggested that the two processes rely on distinct cognitive control mechanisms and possibly different brain circuits.

Pollok et al. (2017, ref. 37) used tDCS approaches to test for causal contributions of dorsolateral premotor cortex to an auditory-motor synchronization and continuation tapping task^37^. Although differences in intensity, depth, focality, and mechanism of stimulation between TMS and tDCS lead us to only cautiously compare the two brain stimulation techniques, both techniques have protocols that down- and up-regulate cortex, and similar perceptual and behavioral effects might be expected to some degree. Pollok et al. (2017, ref. 37) show that both down-regulation and up-regulation to a dorsal premotor target leads to worsening of accuracy in tapping continuation post-metronome, but no change in accuracy during auditory-motor synchronization, suggesting causal involvement of dorsal premotor cortex in precise internal timing of isochronous sequences but not in sensory-guided timing. Specifically, Pollok et al.^37^ show that down-regulation leads to a hastening of tapping with smaller inter-tap intervals, and up-regulation leads to a slowing of tapping with larger inter-tap intervals.

The mechanisms of premotor cortical contributions to timing accuracy for tempo are uncertain, as is how tempo timing relates to predictive beat-based timing in the case of complex rhythms. However, Pollok et al.^37^ show that tDCS down-regulation of dPMC seems to increase, instead of decrease, tendency for negative mean asynchrony, a hallmark of predictive timing, while up-regulation of the area seems to decrease negative mean asynchrony. This is somewhat surprising based on theories of premotor timing prediction, and indicates that the specific mechanisms of dPMC for timing prediction are not yet clear. Pollok et al.^37^ suggest that different cortical areas within motor control networks have distinctive roles in sub-second timing. In support of this hypothesis is dissociation between left posterior parietal cortex^29,40,41^ and left dPMC in specific timing task interference^37^, and the work presented here.

Pollok et al.^37^ used an auditory-motor synchronization and continuation tapping paradigm. We uniquely show here evidence for causal contributions of premotor networks to auditory timing perception in the absence of a motor task, which supports the predictions outlined in the ASAP hypothesis^3^, although it also reveals the need for a more nuanced expansion. Future studies are needed to reveal specifically which premotor networks are involved in different aspects of auditory timing perception.

One limitation of the current design is that our three experiments used adaptive thresholding tests of timing perception acuity that have not been verified to estimate the same perceptual thresholds across tests. That is, the interval discrimination test and the musical tempo and phase subtests of the A-BAT have not yet been shown to be comparable in difficulty and therefore we cannot yet conclude that task ease or difficulty do not contribute to a null result. It is thus not possible to compare the thresholds across the three studies. This concern does not impact positive results. Thus, while our data show evidence for a causal role of dPMC in musical tempo perception, but not in phase perception, this cannot be considered a dissociation between tempo and phase timing perception at this time. We can, however, compare like tests across our two studies^29^ and unambiguously state that we observed phase effects with left PPC down-regulation but not with left dPMC down-regulation, and conversely tempo effects with left dPMC down-regulation but not with left PPC down-regulation. Further work, with explicitly matched difficulty across perceptual tests will be needed to dissociate between different types of timing perception.

Another limitation of the current approach is that individual differences in perceptual thresholds on these tests combined with known individual differences in response to cTBS protocols^42^ results in considerable variability in the perceptual threshold data. This variability might explain why the linear mixed effects models comparing pre- to post-cTBS change across stimulation condition (sham vs. left dPMC) were not significant, while pre- to post-cTBS threshold means clearly increased after dPMC stimulation, but not sham.

## Conclusion

Findings from the present studies suggest causal involvement of left dPMC in interval timing and musical tempo timing perception, and thus support hypotheses that the motor system plays an active role in timing and beat perception. We found no evidence for causal involvement of the left dPMC in musical phase timing perception. Our studies also demonstrate that tempo, phase, and absolute interval timing might recruit different distributed networks in the brain.

## Materials and Methods

### Participants

Participants were thirty healthy adults (20 female, 10 male), ages 18–34 years (mean = 20.0, SD = 2.92), recruited from the University of California, Merced, student population. All participants were dominantly right-handed and screened for atypical hearing, amusia, and contraindications for TMS including increased seizure risk, unstable medical conditions, metal implants in the body other than dental fillings, neurological or psychiatric illness, history of syncope, and head or spinal cord surgery or abnormalities^35^. Participants were asked to remove all metal jewelry before the TMS treatment. Seven participants reported three or more years of musical training or experience, with an average length of training or experience in this group 11.1 years (SD = 7.86). One participant reported 1 year of musical training or experience. The other twenty-two participants reported no musical training or experience. There were not enough participants with musical training or experience to test whether musical experience modulates the effects of cTBS on left dPMC. The experimental protocol was carried out in accordance with the Declaration of Helsinki, reviewed and approved by the University of California, Merced, Institutional Review Board, and all participants gave informed consent prior to testing.

### Interval Timing Discrimination test

An adaptive test of absolute interval timing was used to determine a psychoacoustic threshold for detecting differences in timing between two auditory stimuli. This was a single-interval duration discrimination test, similar to that used by Grube et al. (2010, ref. 36) and implemented in MathWorks’ MATLAB (Natick, MA) using custom-designed functions and the Psychophysics Toolbox, Version 3. Perceptual threshold from the interval timing perception task was used to represent perceptual acuity for sub-second interval discrimination. An increase in threshold can be interpreted as a decrease in perceptual acuity. Specifically, this threshold indicates the minimum interval duration difference that cannot be correctly identified as different. Interfering with normal activity in timing networks involved in this timing task would be expected to raise the perceptual threshold determined by this test. Stimulus beeps were created using MATLAB and were 200 Hz pure tones that lasted 0.1 sec each. Each participant performed the test before and immediately after application of cTBS. In this single-interval duration discrimination test, participants were instructed to make a “same” or “ different”judgment between a reference interval of variable duration, presented first, and a target interval, presented second, for 50 trials. Intervals refer to the duration of silence between pairs of tones; reference intervals were 300, 360, 420, 480, 560, and 600 milliseconds presented in a randomized order. The initial target interval duration was 90% of the reference interval, and was adaptively decreased by 6% or increased by 12% after every two consecutive correct or one incorrect response, respectively. Discrimination thresholds (as a percentage) were calculated as the mean of the absolute value of the difference between the target and reference interval of the last six incorrect trials. The adaptive method we used was a combined transformed and weighted method, using the 1-up 2-down method^43^ with asymmetric step sizes^20^ S_up_ = 2S_down_. We propose the equilibrium point is described by S_down_P(DOWN) = S_up_[1-P(DOWN)], where P(DOWN) = [P(X_p_)]^2^ as in Levitt (1971, ref. 43). Solving for the convergence point P(X_p_) gives √2/3 = 0.816, meaning this procedure estimates the interval length for which a correct discrimination would be given 81.6% of the time. See ref. 29 for more details about the stimuli and adaptive procedure on this test.

#### Beat-Based Timing Test

The Beat Alignment Test, version 2 (BAT)^45^ was designed to test beat perception in a purely perceptual manner that does not require rhythmic movement usually used to assess beat perception. Musical excerpts are presented with an added beep track that is either on-beat, with beeps corresponding to the beat, or perturbed with a tempo or phase manipulation. Each participant performed an adaptive version of each of the BAT subtests (A-BAT IBI and A-BAT PHA, described below) before and after application of cTBS. An increase in threshold on these subtests can be interpreted as a decrease in perceptual acuity for detecting timing differences between music and the beep track. Specifically, these thresholds indicate the minimum tempo or phase differences that cannot be correctly identified as different. Interfering with normal activity in timing networks involved in tempo timing or phase timing would be expected to raise perceptual thresholds determined by these tests.

#### Tempo Timing Perception test

The A-BAT IBI^29,45^ is an adaptive version of the inter-beat interval (IBI) subtest of the BAT^45^. It is a test of tempo timing perception with musical stimuli that adapts in difficulty based on participant performance and determines beat-based timing thresholds for inter-beat interval changes. Musical excerpts are presented with a beep track that is either on-beat, with beeps corresponding to the beat, or perturbed with a tempo manipulation. Participants were instructed to discriminate between correct and altered IBIs in 26 trials by responding after hearing the musical excerpt by button press in a forced-choice task (response alternatives: on-beat or off-beat). See ref. 29 for more details about the stimuli and adaptive procedures.

#### Phase Timing Perception test

The A-BAT PHA^29,45^ is an adaptive version of the phase (PHA) subtest of the BAT^45^. It is a test of phase timing perception with musical stimuli that adapts in difficulty based on participant performance and determines beat-based timing thresholds for detecting shifts in phase. Musical excerpts are presented with an added beep track that is either on-beat, with beeps corresponding to the beat, or perturbed with a phase shift manipulation. Participants were instructed to discriminate between correct and altered phase in 26 trials by responding after hearing the musical excerpt by button press in a forced-choice task (response alternatives: on-beat or off-beat). See ref. 29 for more details about the stimuli and adaptive procedures.

### TMS

cTBS (described by ref. 35), was applied to down-regulate cortical activity at left dPMC or in a sham stimulation condition. The protocol used was a 40-sec train of three pulses at 50 Hz, repeated at 200-millisecond intervals, for a total of 600 pulses^35^. This cTBS protocol was applied at 80% of the participant’ s active motor threshold (AMT), while adhering to safety guidelines for participants and the equipment. If a participant’s 80% of AMT was a greater intensity than can safely be administered with our system, we stimulated at the maximum intensity that was safe. AMT was determined for each participant as the lowest stimulator intensity sufficient to produce a visible twitch with single pulse TMS to left motor cortex in 5 of 10 trials in the first dorsal interosseous (FDI) muscle of the right hand during isometric contraction. Although visible twitch was used to determine AMT, the best location in left motor cortex for right FDI activation was determined by comparing motor-evoked potentials’ size and consistency. Motor-evoked potentials were recorded when at rest, with Ag/AgCl sintered electrodes placed over the belly of the FDI muscle with a ground electrode placed over bone near the right elbow. For single-pulse TMS to primary motor cortex, the figure of eight coil (Magstim, D702 double 70 mm coil, Carmarthenshire, United Kingdom) was placed tangential to the head at an angle of ∼ 45° from the anterior–posterior midline^46^. After AMT was determined, cTBS was applied to left dPMC (experimental condition) or left M1 with the coil facing away from the participant’s head (sham stimulation condition). All participants received both stimulation conditions, in a randomized order, with a minimum of 7 days between each condition.

### Neuronavigation

Brain stimulation was guided using the Magstim Visor 2 3-D motion capture neuronavigation system. The system enabled scaling the Talairach brain using individual participant’s head size and shape. We used 3-D coordinates determined from previous literature for the left dPMC target site, determined using an activation likelihood meta-analysis of 43 imaging studies, reported by Chauvigne, Gitau, & Brown (2014, ref. 38). See Figure 1 for coil placement and orientation^46^.

### Data analysis

All perceptual thresholds were determined using the above described adaptive perceptual tests.

### Statistics

Changes pre- to post-cTBS in perceptual acuity (i.e. perceptual threshold) were analyzed with IBM© SPSS© Statistics, Version 20, using paired samples t-tests for each of the stimulation conditions (dPMC and sham). Additionally, we used linear mixed effects models created in R 3.3.2^47^, using the *lmer* function from the lme4 package (version 1.1.13) with a fixed effect for pre- versus post-cTBS and random effects for condition and for participant. P values were obtained by likelihood ratio tests of the full model with the effect in question against the model without the effect in question^39^.

## Additional Information

### Data availability

All datasets from the current experiments are available upon reasonable request from the corresponding author.

### Author Contributions

J.M.R.: Conception and design of the experiment, collection, analysis and interpretation of the data, designing figures, drafting and revision of the manuscript.

J.R.I.: Conception and design of the experiment, interpretation of the data, drafting and revision of the manuscript.

R.B.: Conception and design of the experiment, interpretation of the data, drafting and revision of the manuscript.

